# Multiple rotavirus species encode fusion-associated small transmembrane (FAST) proteins with cell type-specific activity

**DOI:** 10.1101/2023.04.07.536061

**Authors:** Vanesa Veletanlic, Kylie Sartalamacchia, Julia R. Diller, Kristen M. Ogden

**Affiliations:** Department of Pediatrics, Vanderbilt University Medical Center, Nashville, Tennessee, USA; Department of Pathology, Microbiology, and Immunology, Vanderbilt University Medical Center, Nashville, Tennessee, USA

## Abstract

Fusion-associated small transmembrane (FAST) proteins are viral nonstructural proteins that mediate cell-cell fusion to form multinucleated syncytia. We previously reported that human species B rotavirus NSP1-1 is a FAST protein that induces syncytia in primate epithelial cells but not rodent fibroblasts. We hypothesized that the NSP1-1 proteins of other rotavirus species could also mediate cell-cell fusion and that fusion activity might be limited to cell types derived from homologous hosts. To test this hypothesis, we predicted the structure and domain organization of NSP1-1 proteins of species B rotavirus from a human, goat, and pig, species G rotavirus from a pigeon and turkey, and species I rotavirus from a dog and cat. We cloned these sequences into plasmids and transiently expressed the NSP1-1 proteins in avian, canine, hamster, human, porcine, and simian cells. Regardless of host origin of the virus, each NSP1-1 protein induced syncytia in primate cells, while few induced syncytia in other cell types. To identify the domains that determined cell-specific fusion activity for human species B rotavirus NSP1-1, we engineered chimeric proteins containing domain exchanges with the p10 FAST protein from Nelson Bay orthoreovirus. Using the chimeric proteins, we found that the N-terminal and transmembrane domains determined the cell type specificity of fusion activity. Although the species and cell type criteria for fusion activity remain unclear, these findings suggest that rotavirus species B, G, and I NSP1-1 are functional FAST proteins whose N termini play a role in specifying the cells in which they mediate syncytia formation.

**IMPORTANCE:** Mechanisms of membrane fusion and determinants of host range for pathogens remain poorly understood. Improved understanding of these concepts could open new areas for therapeutic development and shed light on virus epidemiology. Our analyses of NSP1-1 proteins from species B, G, and I rotaviruses provide insights into the diversity of domain features tolerated by functional FAST proteins. Further, the observation that all putative FAST proteins tested can induce syncytia formation in at least some cell types provides evidence that rotaviruses that encode NSP1-1 proteins are fusogenic viruses. Finally, although the criteria for their specificity remain unclear, our observations regarding fusion capacities of different NSP1-1 proteins and of chimeric FAST proteins suggest a potential role for rotavirus FAST proteins in determining the efficiency of viral replication within a given host or cell type.

## INTRODUCTION

Determinants of tropism and pathogenesis are incompletely understood for many viruses, including rotavirus, an important cause of diarrheal disease (1). Rotavirus is a member of the order Reovirales, which contains viruses with segmented double-stranded (ds) RNA genomes (2, 3). Rotaviruses are classified into nine species, rotavirus species A (RVA) through RVD and RVF through RVJ, which can be further resolved into two major clades (https://talk.ictvonline.org/) (4–6). RVA, RVC, RVD, and RVF form clade 1, while RVB and RVG through RVJ form clade 2. Rotaviruses exhibit limited host range and cell tropism, but aside from receptor-binding specificity, the molecular bases of these restrictions are unknown, particularly for rotavirus species other than RVA (7, 8). Most human rotavirus diarrheal disease is caused by RVA and affects infants and young children (2). RVA also causes diarrheal disease in other mammals and in birds. In many cases, specific RVA genotypes are associated with infection of a given animal host, though there is evidence of interspecies transmission and reassortment events (9–15). RVB, RVC, RVH, and RVI have been detected in domesticated mammals, whereas RVD, RVF, and RVG have been detected only in birds (16, 17). RVJ has been detected in bats (18). While RVB is more commonly detected in diarrheic pigs (19–21), it has been associated with sporadic outbreaks of diarrheal disease in humans (22–25). Although the symptoms of RVB gastroenteritis resemble those of RVA, RVB more often causes disease in adults than in infants and young children (26, 27). Sequence analyses suggest the RVBs affecting humans are distinct from those affecting other animals (28, 29); thus, sources of RVB epidemics and reasons these viruses primarily cause disease in adults are unknown. In many cases it remains unclear why some rotavirus species or strains cause disease in a limited range of hosts or in hosts of specific ages.

In contrast to the other ten segments of its dsRNA genome, the predicted gene organization and functions of encoded proteins for the NSP1 segment differ between the two rotavirus clades (30–33). For RVA, the NSP1 segment encodes well characterized innate immune antagonist protein NSP1. RVA NSP1 clusters phylogenetically according to host species, which suggests a potential role in host range restriction (34, 35). For RVB, RVG, and RVI, the NSP1 segment contains two overlapping open reading frames (ORFs) whose encoded products have little predicted homology with known proteins (36). Both encoded proteins are highly divergent among rotavirus species (∼15%-40% identity) and more conserved within a given species (∼40%-100% identity) (unpublished observations). The smaller ORF encodes NSP1-1, which is about 100 amino acids long (36, 37). We previously published evidence indicating that human RVB (HuRVB) NSP1-1 is a fusion-associated small transmembrane (FAST) protein, and we hypothesized that it may play a role in cell tropism (33). Based on sequence analysis, we predicted that RVG and RVI NSP1-1 also may be FAST proteins, but their function has not been directly tested.

Viral FAST proteins are small (∼90-200 amino acids), plasma membrane-spanning proteins that mediate cell-cell fusion at neutral pH and without a specific trigger, resulting in the formation of multinucleated syncytia (reviewed in (38–40)). Unlike the fusion proteins of enveloped viruses, FAST proteins are nonstructural proteins expressed during infection. FAST proteins have been identified not only in HuRVB, but in the genomes of several orthoreoviruses and aquareoviruses (reoviruses), which are also members of the order Reovirales (3, 33, 39, 40). Most current knowledge of FAST protein domain organization and function comes from studies of reovirus FAST proteins, and it is unclear whether rotavirus FAST proteins differ in features or mechanism.

Reovirus FAST proteins have little sequence similarity, but each is composed of a short N-terminal ectodomain, a central transmembrane (TM) domain, and a longer C-terminal endodomain (39, 40). Reovirus FAST proteins are acylated, often at the N terminus but sometimes just after the TM domain, and the endodomain contains a juxtamembrane polybasic region and a predicted amphipathic helix. Often, the three FAST proteins domains can be functionally interchanged (41–44). Reovirus FAST proteins form multimeric complexes at the plasma membrane (43, 45). They interact with the lipid bilayer of closely apposed cells through hydrophobic residues and/or a fatty acid modification in the N terminus (46–51). These interactions are proposed to favor lipid mixing, creating a state that can favor progression to the fusion pore (39, 40). The amphipathic helix in the reovirus FAST protein endodomain is thought to partition into the curved membrane of the pore on the inner leaflet and stabilize it. Then, cellular proteins promote pore expansion and syncytia formation (52–54).

It is possible that FAST proteins contribute to cell type-tropism. While reovirus FAST proteins are not thought to bind specific host cell receptors, in some cases host molecules that interact with the C terminal endodomain and differ among FAST proteins have been identified (52–54). HuRVB NSP1-1 expression results in syncytia formation in primate (human or African green monkey) epithelial cells but not in rodent (hamster or mouse) fibroblasts (33). This observation suggests the possibility of host-or cell type-specific interactions with NSP1-1 that mediate cell-cell fusion. FAST protein expression enhances viral replication in cultured cells (33, 55, 56) and could conceivably contribute to rotavirus replication efficiency in a specific host or tissue during natural infection. In the current study, we sought to predict structural and functional features of NSP1-1 proteins from species B, G, and I rotaviruses derived from different host animals, determine whether they are FAST proteins and can mediate cell-cell fusion efficiently in cell lines derived from different tissues or animal hosts, and identify the protein domains that dictate cell type-specific fusion activity. Results of these studies provide insights into the diversity of features of FAST proteins and identify FAST protein domains that influence cell type-specific activity.

## RESULTS

### Rotavirus NSP1-1 proteins are predicted to share features

Using a variety of algorithms (57–62), we aligned and predicted sequence and structural motifs in RVB, RVG, and RVI NSP1-1. In addition to HuRVB NSP1-1, we included NSP1-1 from pig (porcine; Po) and goat (caprine; Cp) RVB, which also cause diarrhea in their hosts (63, 64). We also included NSP1-1 sequences from pigeon (avian; Av) RVG and turkey (gallinaceous; Ga) RVG (65). RVG has been rarely associated with runting and stunting syndrome in chickens and turkeys (17, 66). Finally, we included NSP1-1 sequences from canine (Ca) RVI, which was sequenced from sheltered dogs, and feline (Fe) RVI, which was sequenced from a diarrheic cat (67, 68). An N-myristoylation site was predicted at amino acids two through seven for every complete RVB, RVG, and RVI NSP1-1 sequence in GenBank (**Fig. 1A-B**) ((33) and not shown). TM helices were identified in RVB, RVG, and RVI NSP1-1 sequences, with the N terminus predicted to be extracellular and the C terminus cytoplasmic. Each NSP1-1 contains multiple basic residues C-terminal to the predicted TM domain. For analyzed RVB, RVG, and RVI NSP1-1 sequences, residues preceding and following the TM domain were predicted to form helices. For PoRVB and CpRVB NSP1-1, the endodomain helix is predicted to be amphipathic (**Fig. 1C**). These motifs suggest a model of RVB NSP1-1 in which a myristoylated extracellular N-terminal ectodomain, which may interact with lipids, precedes a TM domain and a cytoplasmic endodomain containing a polybasic region that, in some cases, is in an amphipathic helix and may interact with the plasma membrane inner leaflet (**Fig. 1B**). Similar topology is predicted for RVG and RVI NSP1-1, though the endodomains are smaller, and no amphipathic helices were modeled.

**Figure 1.**
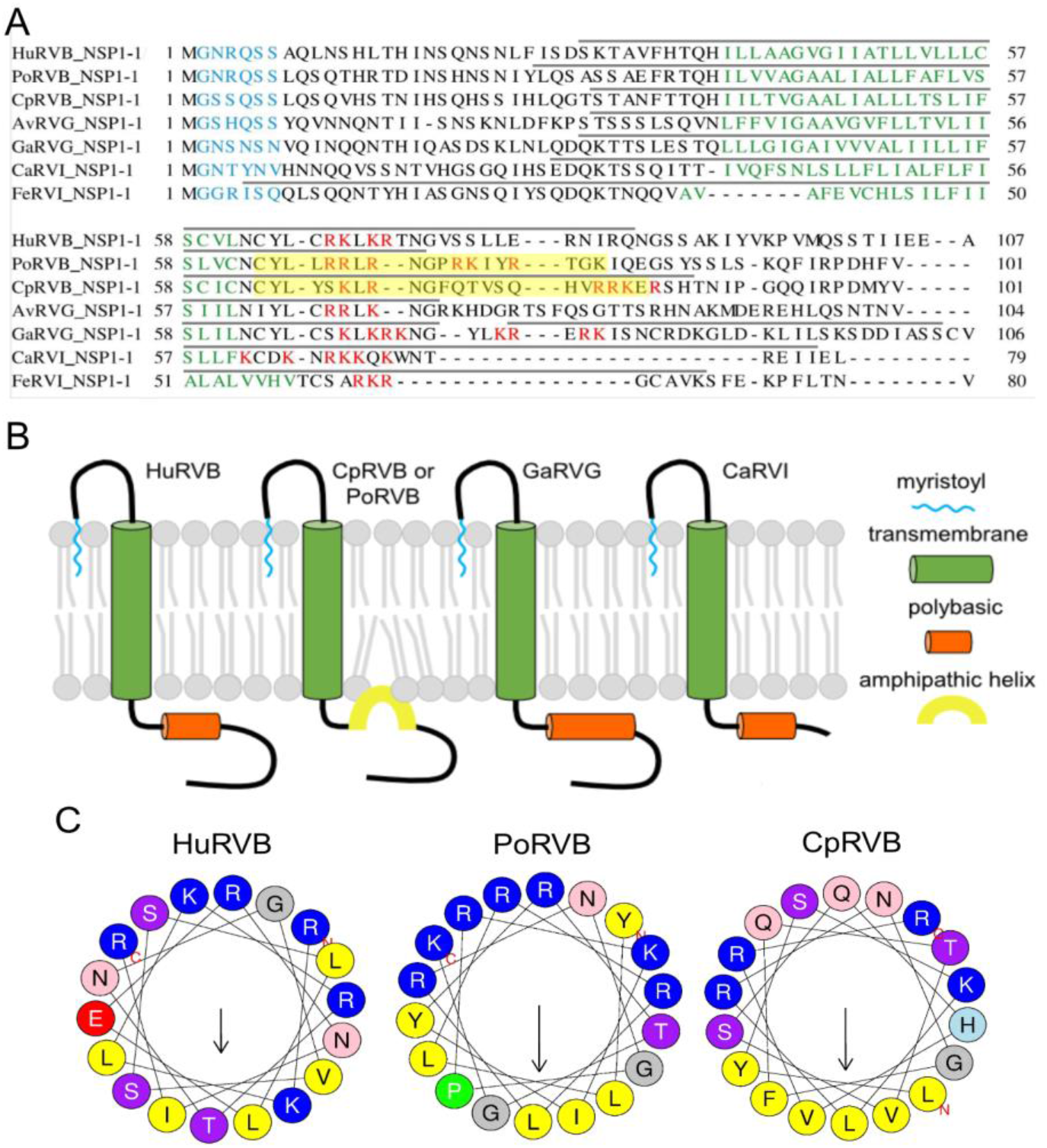
Sequence alignments (A) and cartoon model highlighting predicted features (B) of NSP1-1 from RVB, RVG, and RVI. Predicted N-myristoylation motifs (cyan), transmembrane helices (green), polybasic regions (orange), and amphipathic helices (yellow) are indicated (57, 58, 60, 62). Gray lines above sequences in (A) indicate residues predicted to fold into helices (59). (C) Helical wheel diagram for an 18-amino acid window in the predicted helical region of Hu, Po, or Cp RVB NSP1-1 following the TM domain. Hydrophobic residues (yellow) cluster opposite basic (dark blue) and uncharged polar (pink and purple) residues for PoRVB and CpRVB NSP1-1, but not for HuRVB NSP1-1 (57).

### RVB, RVG, and RVI NSP1-1 mediate syncytia formation in human cells

To test the hypothesis that all rotavirus NSP1-1 proteins are FAST proteins that mediate cell-cell fusion, we transfected human embryonic kidney 293T cells with vector alone or plasmids encoding HuRVB, PoRVB, CpRVB, AvRVG, GaRVG, CaRVI, or FeRVI NSP1-1 (37, 63–65, 67, 68). Then, we examined the appearance of the cell monolayer and organization of nuclei and F-actin. F-actin tends to cluster near the cell periphery but forms a network throughout the cell (69), and syncytia contain multiple nulcei. While vector-transfected cells were indistinguishable from mock-transfected cells, transfection with any NSP1-1 expression plasmid changed morphology from distinct cells to a monolayer pockmarked by smooth round or oval-shaped syncytia lacking defined cell edges (**Fig. 2A**). Multiple nuclei were often clustered near the center or at the edges of these syncytia, and while actin was detectable in these areas, actin staining was noticeably lighter and had a different pattern. Combined with their predicted domain features, these observations suggest that RVB, RVG, and RVI NSP1-1 from rotaviruses that infect different animals are FAST proteins that can mediate cell-cell fusion. Average diameters of syncytia formed by GaRVG, CaRVI, and FeRVI were smaller than those formed by HuRVB NSP1-1 (**Fig. 2B**), and they were detected less frequently, which may suggest differences in the nature or efficiency of interactions in human cells.

**Figure 2.**
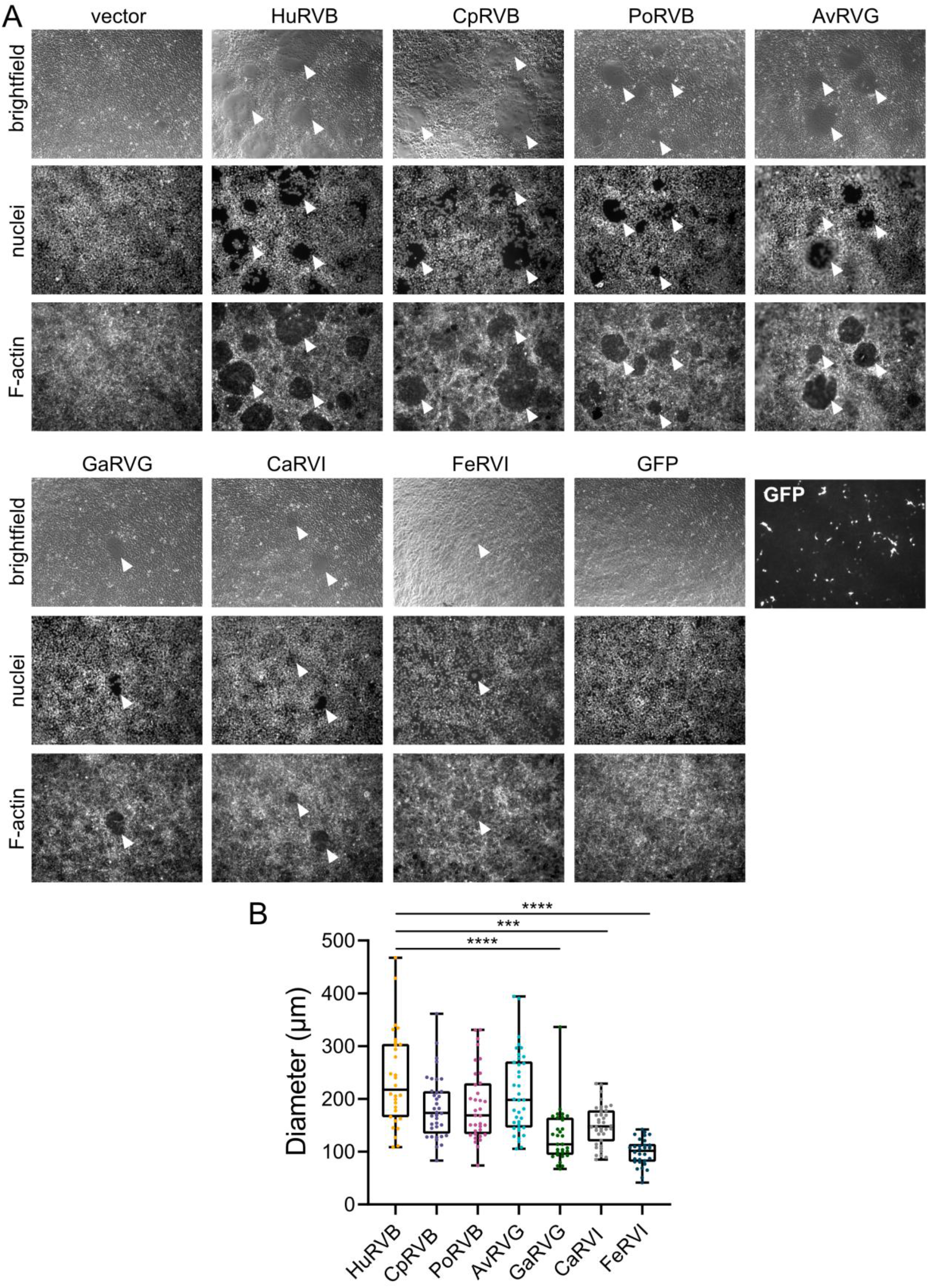
RVB, RVG, and RVI encode functional FAST proteins. (A) Brightfield and DAPI or rhodamine-stained images of 293T cells transfected with pCAGGS alone (vector) or pCAGGS expressing HuRVB, CpRVB, PoRVB, AvRVG, GaRVG, CaRVI, or FeRVI NSP1-1 or GFP at 18 h post transfection. White arrowheads indicate syncytia. (B) Bar graph showing diameters of syncytia for HuRVB, CaRVB, PoRVB, AvRVG, GaRVG, CaRVI, and FeRVI NSP1-1 at 18 h post transfection in 293T cells. *n* ≥ 30 syncytia. ***, P < 0.001; ****, P < 0.0001 compared with HuRVB NSP1-1 diameter by Kruskal-Wallis test with Dunn’s multiple comparisons.

### Some rotavirus NSP1-1 proteins with a C-terminal peptide tag are fusion active

To enable detection of HuRVB NSP1-1, we previously engineered a FLAG peptide at the N or C terminus and found that N-terminally tagged FLAG-NSP1-1 was expressed in individual 293T cells with distinct edges, while C-terminally tagged NSP1-1-FLAG was expressed in syncytia (**Fig. 3A)** (33). These findings are consistent with disruption of a myristoyl moiety on the N terminus of HuRVB NSP1-1 by addition the FLAG peptide (**Fig. 1A-B**) and suggested that a free C terminus was not necessary for fusion activity. Since all NSP1-1 proteins are predicted to be N-terminally myristoylated, it is likely that addition of a FLAG peptide would ablate cell-cell fusion activity for all these FAST proteins, as it did for HuRVB NSP1-1. To enable detection and determine the requirement for a free C terminus for NSP1-1 proteins other than HuRVB NSP1-1, we engineered a FLAG peptide at the C terminus for our panel of RVB, RVG, or RVI NSP1-1 proteins. While FLAG-tagged AvRVG and RVI NSP1-1 proteins were undetectably expressed or mislocalized, all RVB NSP1-1 proteins and GaRVG NSP1-1 were detected and could mediate cell-cell fusion in 293T cells (**Fig. 3A**). In each case, there was a trend towards smaller syncytia diameter induction for the FLAG-tagged compared with the untagged form of the protein, with statistical significance for CaRVB, PoRVB, and GaRVG NSP1-1 (**Fig. 3B**). These findings suggest potential differences in functional interactions of the NSP1-1 endodomain among rotavirus species and strains, with a critical role for this domain for RVI and AvRVG NSP1-1 and a contributing but non-critical role for this domain in fusion for RVB and GaRVG NSP1-1.

**Figure 3.**
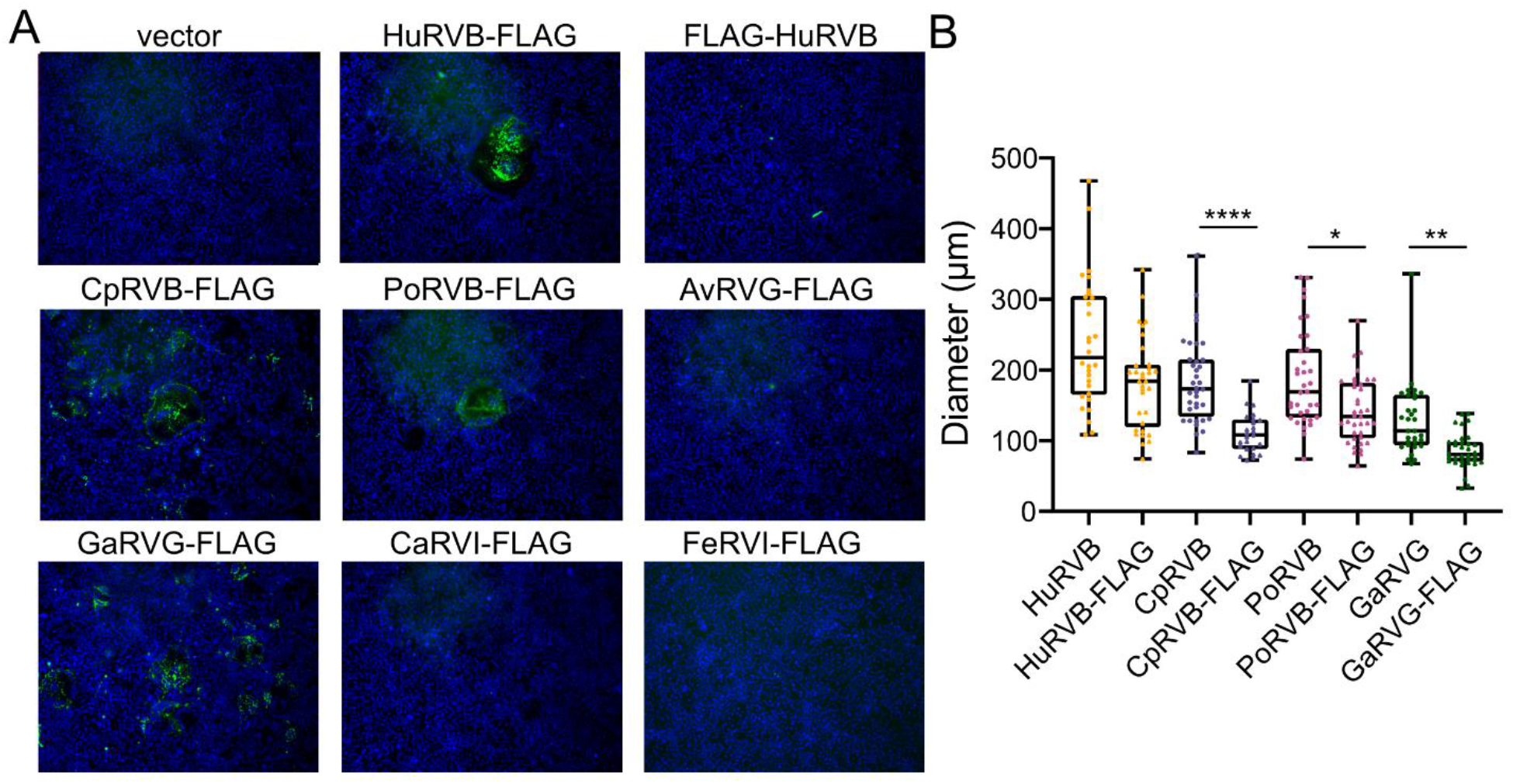
The extreme C terminus is critical for RVG NSP1-1 and RVI NSP1-1 but not RVB NSP1-1 function. (A) DAPI and FLAG-stained images of 293T transfected with pCAGGS alone (vector) or pCAGGS expressing FLAG-HuRVB NSP1-1, HuRVB-NSP1-1-FLAG, CaRVB NSP1-1-FLAG, PoRVB NSP1-1-FLAG, AvRVG NSP1-1-FLAG, GaRVG NSP1-1-FLAG, and FeRVI NSP1-1-FLAG at 18 h. (B) Bar graph showing diameters of syncytia for HuRVB-NSP1-1-FLAG, CaRVB NSP1-1-FLAG, and PoRVB NSP1-1-FLAG at 18 h post transfection in 293T cells. *n* ≥ 30 syncytia. The diameters of FLAG-tagged RVB NSP1-1 are compared to those of untagged RVB NSP1-1 (data duplicated from Fig. 2C). ****, P < 0.0001; **, P < 0.01; *, P < 0.05 by Kruskal-Wallis test with Dunn’s multiple comparisons.

### Rotavirus NSP1-1 proteins exhibit species-specific cell fusion activity

Our prior observation that HuRVB NSP1-1 can induce syncytia formation in primate cells but not in rodent cells raised the possibility that this protein functions as a tropism determinant, promoting virus spread between cells of homologous human or simian but not heterologous rodent hosts (33). To test the hypothesis that NSP1-1 functions in a species-specific manner, we obtained cell lines derived from host animals that are homologous, derived from the same animal species, or heterologous, derived from different animal species than the parent rotavirus from which NSP1-1 sequences were cloned. We transfected them with NSP1-1 expression plasmids and looked for the presence of syncytia. We chose cell lines that are transfectable. Since NSP1-1 proteins form syncytia in human embryonic kidney epithelial cells (**Fig. 2**), in several cases we also used cells of kidney or epithelial cell origin. We transfected cells with the pCAGGS vector alone as a negative control, and we transfected cells with pCAGGS expressing NBV p10 from the Miyazaki-Bali (MB) strain (55, 70, 71) as a positive control, since it induces syncytia formation in at least some cell types that HuRVB NSP1-1 does not (33). In Cos7 cells, an African green monkey kidney fibroblast-like cell line, transfection with each of the RVB, RVG, and RVI NSP1-1 expression plasmids resulted in readily visible syncytia in the cell monolayer, regardless of the animal source of the viral protein (**Fig. 4**). Even CaRVI and FeRVI NSP1-1 proteins, which induced small syncytia in 293T cells (**Fig. 2**), appeared to induce syncytia efficiently in Cos7 cells (**Fig. 4**). In baby hamster kidney BHK-T7 cells, which are heterologous with all parent viruses from which our NSP1-1 sequences were derived, only NBV p10 and AvRVG NSP1-1 proteins induced readily detectable syncytia (**Fig. 5A**). The same was true for porcine kidney epithelial LLC-PK1 cells, for which no syncytia were detected following transfection with PoRVB NSP1-1 (**Fig. 5B**). In chicken embryo fibroblast DF-1 cells, syncytia were detected only following transfection with pCAGGS expressing NBV MB p10, not with any NSP1-1 protein, even those derived from pigeon or turkey RVG (**Fig. 5C**). Finally, in canine kidney fibroblast-like MDCK.1 cells, no differences in cell morphology relative to vector-transfected monolayers were detected for cells transfected with FAST protein-expressing plasmids (**Fig. 5D**). Transfection efficiencies were low in MDCK.1 cells, but we failed to detect syncytia even for NBV MB p10. Together, these observations suggest that NSP1-1 proteins induce syncytia formation in a limited range of cell types, but functional range does not strictly correlate with the animal or tissue origin of a given cell.

**Figure 4.**
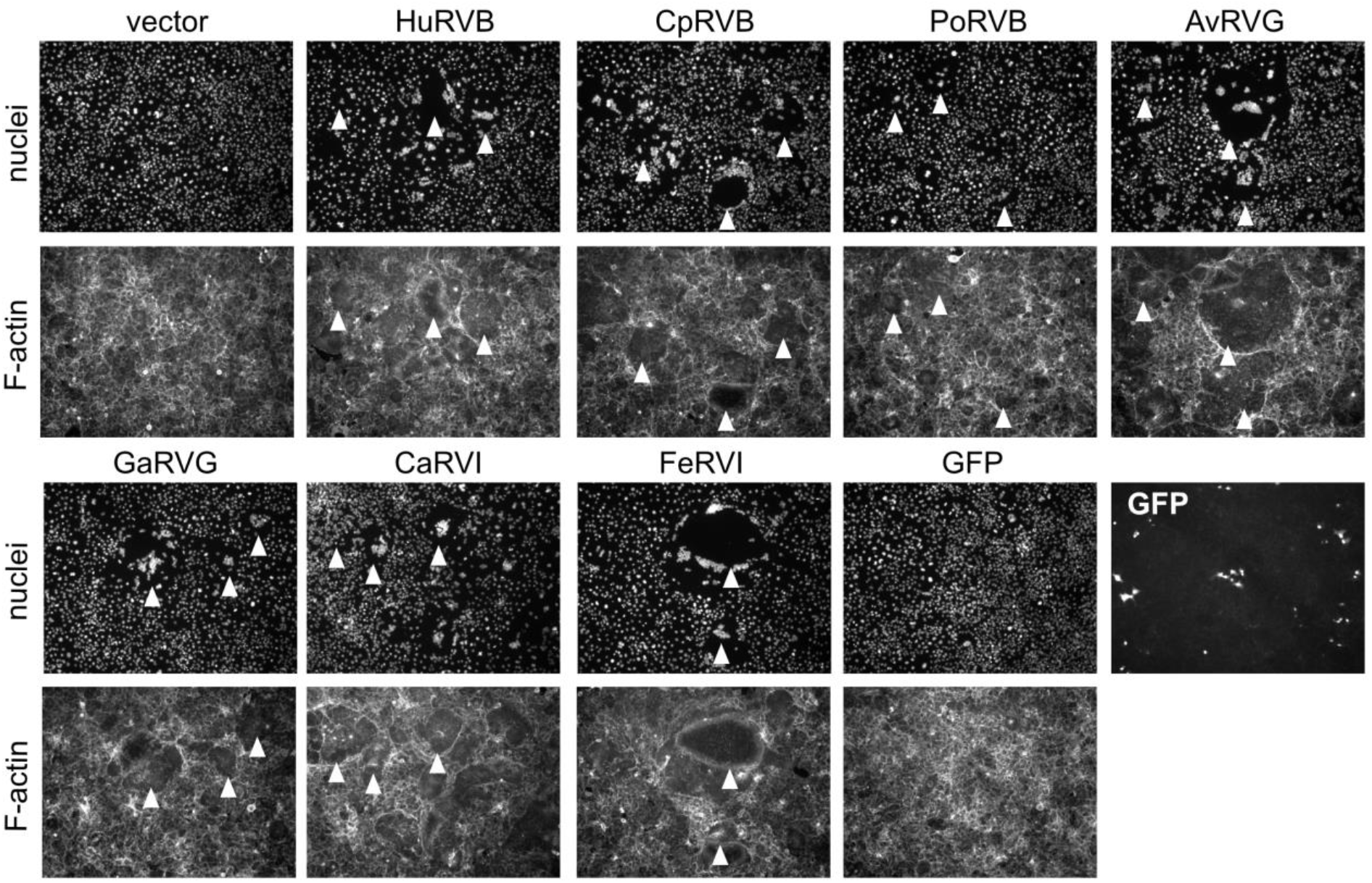
RVB, RVG, and RVI NSP1-1 form syncytia in primate cells. DAPI-stained (nuclei) and rhodamine-stained (F-actin) images of Cos7 cells transfected with pCAGGS alone (vector) or pCAGGS expressing NBV p10, the indicated RVB, RVG, or RVI NSP1-1, or GFP as a transfection control. White arrowheads indicate syncytia.

**Figure 5.**
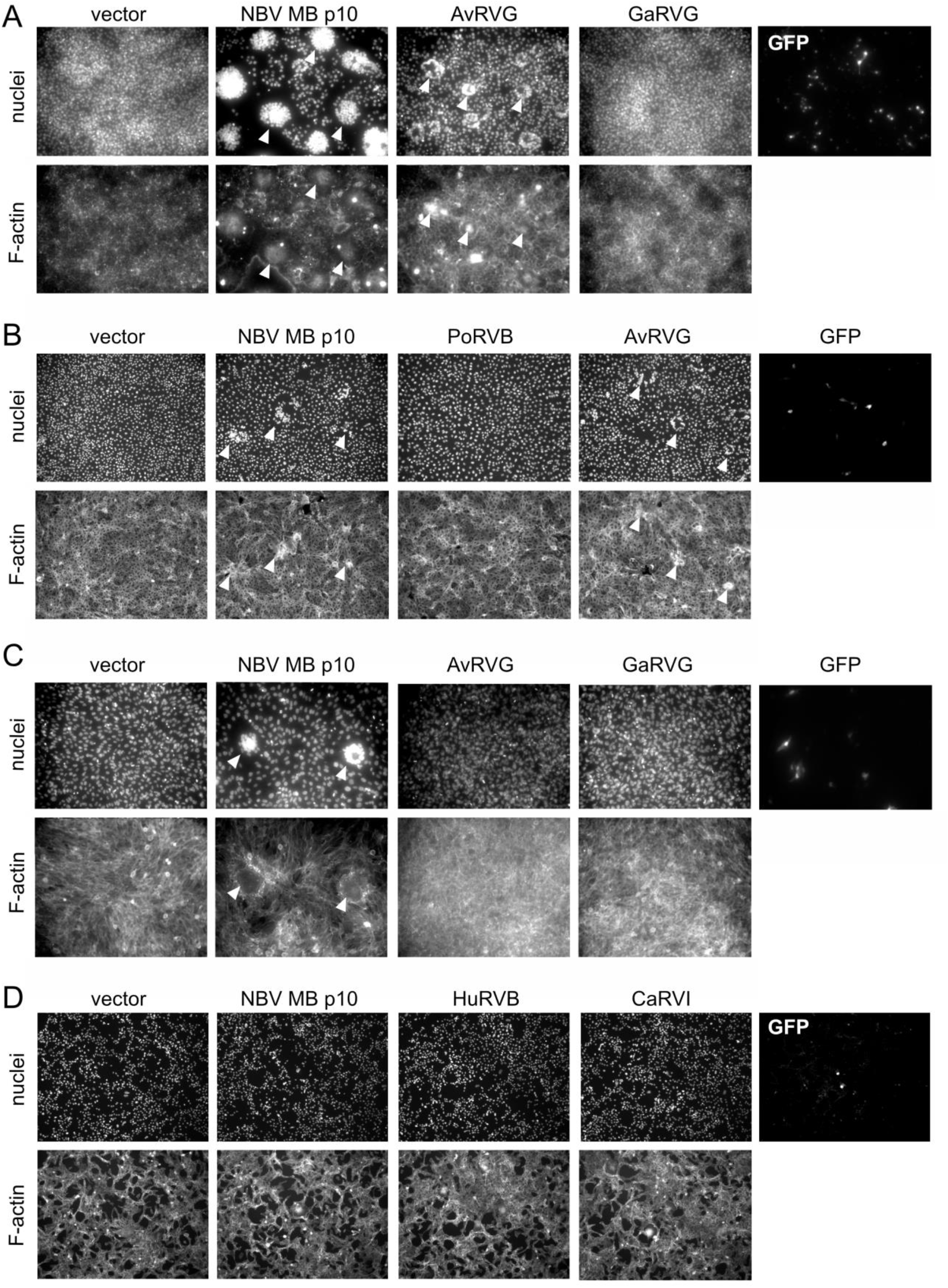
RVB, RVG, and RVI NSP1-1 have limited capacity to form syncytia in epithelial cells from several other animals. DAPI-stained (nuclei) and rhodamine-stained (F-actin) images of BHK (A), LLC-PK1 (B), DF-1 (C), or MDCK.1 (D) cells transfected with vector, NBV p10, GFP, and RVB, RVG, or RVI NSP1-1. Images for selected constructs are shown to represent negative (vector) and positive (NBV p10) controls and the level of transfection (GFP), as well as for selected NSP1-1 proteins. NBVp10 is not a positive control in MDCK.1 cells. White arrowheads indicate syncytia.

### The N terminus can confer FAST protein species specificity

We previously determined that HuRVB NSP1-1 retains fusion activity in human (293T) and simian (MA104) cells but not in rodent (BHK-T7 cells and murine L929 fibroblasts) (33). However, NBV MB p10 induces syncytia when expressed in each of these cell lines. To learn more about FAST protein domain function, we aligned sequences of HuRVB NSP1-1 and NBV MB p10 and engineered chimeric constructs based on the alignments, in which we exchanged N-terminal, TM, and C-terminal domains between the two FAST proteins (**Fig. 6A-B**). NBV MB p10 is palmitoylated at a membrane proximal dicysteine motif C terminal to the TM domain (72); we preserved this motif in our chimeric constructs (**Fig. 6A-B**). We anticipated that all properly folded constructs should induce syncytia formation in 293T cells, since neither parent protein exhibited restricted fusion activity in this cell line. Despite high transfection efficiency, we found that only the chimeric proteins in which the C-termini had been exchanged induced detectable syncytia formation in 293T cells (**Fig. 6C**). This finding suggests that these chimeric proteins are properly folded and post-translationally modified, whereas those with N-terminal or TM domain exchanges are not.

**Figure 6.**
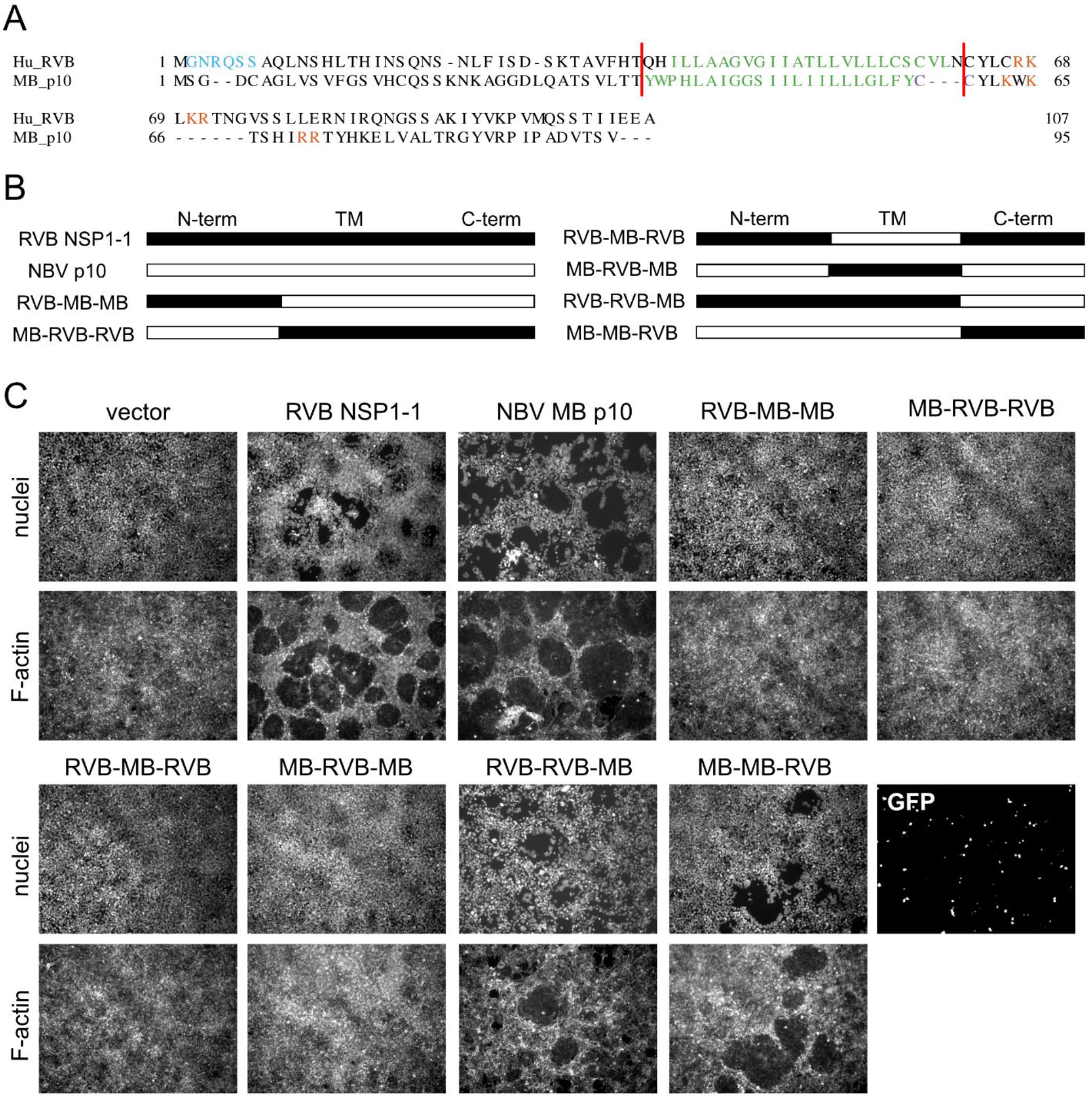
HuRVB NSP1-1 is a modular FAST protein. (A) Alignment of NBV MB p10 and HuRVB NSP1-1. Predicted N-myristoylation motifs (cyan), transmembrane helices (green), polybasic regions (orange), and palmitoylation site (purple) are indicated. Sequences were aligned using MAFFT v7.2. For RVB NSP1-1, features were identified as described in Fig. 1A. For p10, features were described previously (72). (B) Linearized schematic representations of engineered chimeric protein constructs. (C) DAPI-stained (nuclei) and rhodamine-stained (F-actin) images of 293T cells transfected with plasmids expressing HuRVB NSP1-1, NBV MB p10, the indicated RVB/MB chimeric protein, or GFP as a control for transfection efficiency.

To identify the FAST protein domain that confers species or cell-type specificity, we transfected BHK-T7 cells with plasmids encoding parental HuRVB NSP1-1 and NBV MB p10 proteins or with the chimeric C-terminally exchanged proteins. As previously observed (33), RVB NSP1-1 failed to induce syncytia formation in BHK-T7 cells, even when transfection efficiency was quite high (**Fig. 7**). Only when the N-terminal and TM domains of NBV p10 were present did we detect syncytia in the BHK-T7 monolayer. The chimeric protein containing the N-terminal ectodomain and TM domain of RVB NSP1-1 and C-terminal endodomain of NBV p10 induced no detectable change in monolayer appearance. Although the C-terminal endodomain is thought to interact with host proteins to mediate pore formation for some reovirus FAST proteins (52–54), our findings suggest that the N terminus can contribute to FAST protein species-specific activity.

**Figure 7.**
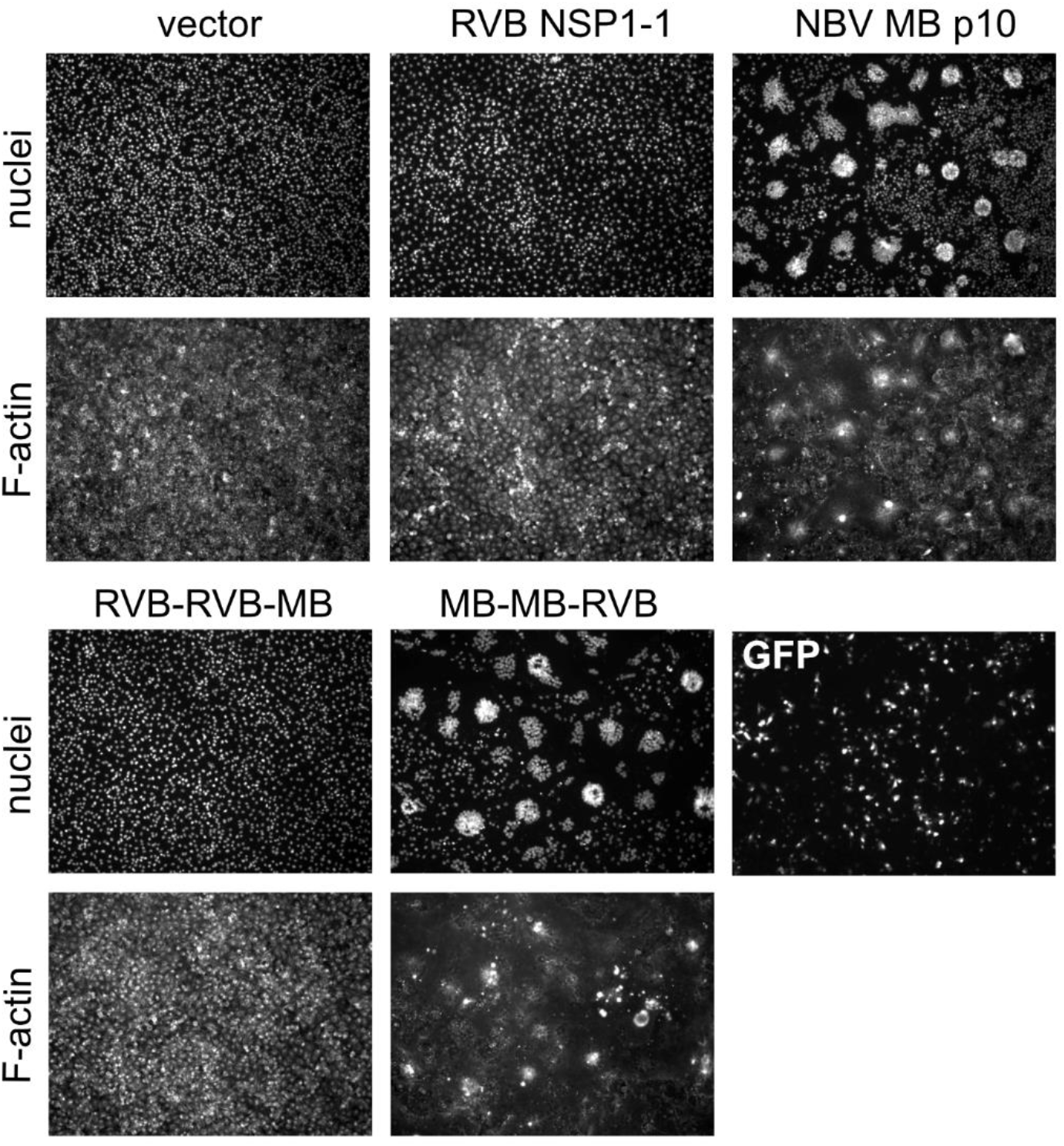
The N-terminal and transmembrane domains determine species-specific fusion activity for RVB NSP1-1 and NBV p10. DAPI-stained (nuclei) and rhodamine-stained (F-actin) images of BHK-T7 cells transfected with plasmids expressing HuRVB NSP1-1, NBV MB p10, the indicated RVB/MB chimeric protein, or GFP as a control for transfection efficiency.

## DISCUSSION

In the current study, we sought to test the hypothesis that NSP1-1 proteins from different rotavirus species are functional FAST proteins. The capacity of all RVB, RVG, and RVI NSP1-1 proteins tested to induce syncytia formation in primate cells suggests that, like orthoreovirus and aquareovirus FAST proteins, they can mediate cell-cell fusion (39, 40) (**Figs. 2 and 4**). Similar to reovirus FAST proteins, RVB, RVG, and RVI NSP1-1 are predicted to be acylated and to contain an N-terminal ectodomain, a central TM domain, and a C-terminal endodomain (39, 40). However, while reovirus proteins are reported to have endodomains that are equal in size or substantially longer than the ectodomains (39), for RVG and RVI NSP1-1 the predicted endodomains are shorter than the ectodomains (**Fig. 1**). While all reovirus FAST proteins described to date are predicted to contain amphipathic alpha helices in the endodomain that splay apart lipid headgroups, lowering the energy barrier to pore formation (39, 40), only a subset of the NSP1-1 proteins we analyzed are predicted to contain such motifs (**Fig. 1**). Thus, some NSP1-1 proteins may employ a distinct mechanism to stabilize pore formation, for example interaction with a cellular protein that contains amphipathic regions and partitions into the inner leaflet of the bilayer. Together, these observations suggest that RVB, RVG, and RVI NSP1-1 are functional FAST proteins, that there is some flexibility in the domain features of FAST proteins, and that FAST proteins of rotaviruses and reoviruses may differ at least modestly in how they induce cell-cell fusion.

We also sought to test the hypothesis that rotavirus NSP1-1 exhibits host species-or cell type-specific activity. This hypothesis was based on the observation that HuRVB NSP1-1 mediated cell-cell fusion in primate epithelial cells but not rodent fibroblasts (33) (**Figs. 6 and 7**). We hypothesized that if NSP1-1 is a host range determinant, then NSP1-1 proteins would mediate syncytia formation efficiently in homologous cells, and they would mediate syncytia formation inefficiently in heterologous cells. Alternatively, if only the tissue type from which a cell line was derived mattered, then NSP1-1 might mediate syncytia formation efficiently in epithelial-derived but not fibroblast-derived cell lines. Unexpectedly, we found that NSP1-1 proteins from RVB, RVG, and RVI from different host animals all mediated readily detectable cell-cell fusion upon expression in primate epithelial (293T) and fibroblast (Cos7) cells (**Figs. 2 and 4**). Rarely did these proteins mediate syncytia formation in other tested cell lines, and when they did, it was not in a homologous cell line. For example, AvRVG NSP1-1 mediated detectable cell-cell fusion in porcine epithelial (LLC-PK1) cells, while PoRVB NSP1-1 did not (**Fig. 4**). These observations suggest that NSP1-1 cell fusion function is limited, but it is not strictly limited to the host in which the virus was initially detected or to cells derived from a specific tissue type. Nonetheless, HuRVB NSP1-1 does mediates cell-cell fusion in primate epithelial but not rodent fibroblast cells, while NBV MB p10 mediates cell-cell fusion in both cell lines (33) (**Figs. 6 and 7**). Our findings indicate that the N-terminal ectodomain, TM domain, or both domains dictate this cell-specific fusion activity, which suggests that these domains participate in specific interactions with host cell molecules (**Figs. 6-7**). Host molecules are required for cell fusion by several reovirus FAST proteins (52–54). While the reptilian orthoreovirus p14 FAST protein endodomain interacts with Grb2 to trigger N-WASP-mediated actin polymerization, aquareovirus p22 FAST protein uses adaptors Intersectin-1 and Cdc42 to trigger N-WASP-mediated branched actin assembly (52, 53). To date, no specific interactions with cellular molecules have been identified for reovirus FAST protein N-terminal ectodomains or TM domains (39, 40). Taken together, our findings indicate that there is cell-specific fusion activity for rotavirus NSP1-1 that is dictated by the N terminus, but they fail to clearly delineate species or cell type criteria for fusion activity.

Why did most NSP1-1 proteins fail to induce syncytia in many tested cell lines? Since we do not have antibodies specific for our panel of NSP1-1 proteins, and many failed to tolerate a peptide tag (**Fig. 3**), NSP1-1 expression in each cell line could not easily be verified. However, several lines of evidence suggest untagged NSP1-1 proteins were expressed. First, detection of GFP expressed from a plasmid indicated that each cell type was transfection competent under the assay conditions (**Fig. 5**). Second, the promoter for the pCAGGS plasmid is functional for high-level protein expression in most cell types. Third, syncytia were formed by p10 in every cell type except MDCK.1, suggesting protein expression was driven from the CAG promoter, and the cells were competent for fusion (**Fig. 5**). Finally, each NSP1-1 construct formed syncytia successfully in 293T and Cos7 cells (**Figs. 2 and 4**). Despite detecting syncytia only in primate cell lines, we think it is unlikely that NSP1-1 mediates syncytia formation only in primates. Indeed, syncytia have been detected in the epithelial cells of the small intestinal villi of rats and pigs infected with RVB (73, 74). It is possible that some rotavirus NSP1-1 proteins lack cell fusion activity *in vivo*, though they likely are maintained in the compact viral genome for a reason. Tight cell-cell apposition, as found between intestinal epithelial cells, may be a requirement for successful NSP1-1-mediated cell-cell fusion. Accordingly, cadherin and other adhesion factors enhance reovirus FAST protein-mediated cell-cell fusion (46). There also may be host factors present or absent in some cell types that promote or restrict syncytia formation. However, most cellular molecules identified to interact with reovirus FAST proteins are broadly expressed, including adaptors in the branched actin assembly network mentioned above and Annexin A1, which interacts with reptilian orthoreovirus FAST protein p14 and promotes fusion pore expansion (52–54). While restriction factors directed towards FAST proteins have not yet been identified, several cellular molecules have been proposed to restrict HIV-1 infection, some of which are expressed in specific cell types (reviewed in (75–78)). Additional studies, perhaps identifying and comparing cellular binding partners of p10 and NSP1-1, may uncover reasons for the specificity of NSP1-1 fusion observed in our experiments.

It is unclear what our observations regarding NSP1-1 behavior in cultured cells suggest about the behavior of NSP1-1 from fusogenic rotaviruses in their natural hosts. FAST proteins may behave differently in the context of a rotavirus during natural infection than when expression is driven by a non-native promoter following plasmid transfection. The lack of tissue culture systems for most of these viruses has limited their study in the laboratory, and whether cell-cell fusion is a contributor to the pathogenesis of each rotavirus from which the NSP1-1 sequences were derived remains unknown (66-68, 79-81). For NBV, FAST protein p10 dramatically increased virus titer and pathogenesis in a mouse model (55). Although NBV exhibits cell type specific replication, it is determined not by p10 FAST, but by the p17 protein, which is encoded on the same segment in a separate open reading frame (82). While only p10 FAST expression is required to permit efficient NBV replication in primate epithelial (Vero) cells, both p10 and p17 expression are required for efficient replication in bat (DemKT1) cells, and another bat homolog of NBV p17 but not homologs from avian or baboon reoviruses could complement this p17 function. Rotaviruses lack a p17 homolog and may have evolved to confer species specificity directly via the FAST protein, or they may employ a different viral protein to confer this property. For the aquareovirus grass carp reovirus, fusion activity of the NS16 FAST protein is enhanced by the expression of another viral protein, NS26, possibly through its interaction with host lysosomes (83, 84). Future studies of fusogenic rotaviruses may reveal new information about cell tropism for these viruses and whether additional viral proteins modulate tropism or FAST protein activity.

In summary, our observations provide evidence that, although their predicted features deviate in some cases from those of reovirus FAST proteins, the NSP1-1 proteins of species B, G, and I rotaviruses are FAST proteins that can mediate syncytia formation in at least some cell types. The N terminus of HuRVB NSP1-1 influences the cell type specificity of its fusion activity. Many questions and much work remain to be done to understand the biological mechanism and function of NSP1-1 in the context of a rotavirus and its natural host and to elucidate determinants of rotavirus host range and cell tropism.

## MATERIALS AND METHODS

### NSP1-1 Alignment and Prediction of Protein Features

NSP1-1 sequences were obtained from GenBank. Accession numbers for NSP1-1 FAST sequences are ADF57900 (HuRBV), ASN74338 (PoRVB), ASV45172 (CpRVB), AXF38051 (AvRVG), ASV45159 (GaRVG), YP_009130668 (CaRVI), and AQX34665 (FeRVI). The accession number for the Nelson Bay orthoreovirus Miyazaki-Bali strain p10 is BAT21545. For Figs. 1A and 6A, amino acid sequences were aligned using MAFFT v7.2 using the E-INS-I strategy (62). For NSP1-1 protein feature prediction, myristoylation motifs were identified using ExPasy Scan Prosite (61). Transmembrane sequences were identified using DeepTMHMM (60). Amphipathic helices were identified using Proteus2 and heliQuest (57, 58). Sequences predicted to fold into helices were identified using AlphaFold2 through ColabFold (59). Model #1 was used for predictions of helical regions shown in **Fig. 1A**. In most cases, all models for a given NSP1-1 sequence were similar. The TM domain, polybasic region, and palmitoylated cysteines in NBV p10 were identified previously (72).

### Plasmids

NBV (Miyazaki-Bali) p10 in pCAGGS has been described previously (56). HuRVB (Bang117) NSP1-1 in pCAGGS and N-and C-terminally FLAG-tagged forms of this construct have been described previously (33). pLIC6 was constructed by engineering a ligation-independent cloning site into mammalian expression plasmid pCAGGS. Sequences encoding CpRVB NSP1-1, PoRVB NSP1-1, AvRVG NSP1-1, GaRVG NSP1-1, CaRVI NSP1-1, and FeRVI NSP1-1 with a C-terminal FLAG peptide inserted prior to the STOP codon were synthesized (GenScript). Ligation-independent cloning following PCR amplification with appropriate primers and T4 DNA polymerase treatment was used to clone the sequences either without (untagged) or with (tagged) the C-terminal FLAG peptide into pLIC6. Sequences encoding chimeric HuRVB NSP1-1 and NBV MB p10 proteins with exchanged N termini, TM domains, and C termini were synthesized (GenScript). Ligation-independent cloning following PCR amplification with appropriate primers and T4 DNA polymerase treatment was used to clone the sequences into pLIC6. Nucleotide sequences of plasmid constructs were verified by Sanger sequencing.

### Cells

Human embryonic kidney 293T cells were grown in Dulbecco’s modified Eagle’s minimal essential medium (DMEM) (Corning) supplemented to contain 10% fetal bovine serum (FBS) (Gibco). Monkey kidney fibroblast Cos7 cells were grown in DMEM supplemented to contain 10% FBS. Baby hamster kidney cells expressing T7 RNA polymerase under control of a cytomegalovirus promoter (BHK-T7) (85) were grown in DMEM supplemented to contain 5% FBS, 10% tryptose phosphate broth (Invitrogen), and 1% nonessential amino acids (Corning), with 1 mg/ml G418 (Invitrogen) added during alternate passages. Canine kidney fibroblast-like MDCK.1 cells were grown in Eagle’s minimal essential medium (Corning) supplemented to contain 10% FBS. Porcine kidney epithelial LLC-PK1 cells were grown in Medium 199 with Earle’s salts (Gibco) plus 2.2 g/L sodium bicarbonate and supplemented to contain 3% FBS. Chicken embryo fibroblast UMNSAH/DF-1 cells were grown in DMEM supplemented to contain 10% FBS. All culture media were supplemented to contain 2 mM L-glutamine (Corning). Except for BHK-T cells, culture media also contained 100 units/mL penicillin and 100 µg/mL streptomycin (Corning).

### Antibodies

Monoclonal mouse anti-FLAG antibody (Sigma), Alexa Fluor 488-conjugated anti-mouse IgG (Invitrogen), and rhodamine phalloidin (Invitrogen) are commercially available.

### Cell Transfection and Imaging

For differential interference contrast imaging, 293T cells (∼2 × 10^5^ per well) in 24-well plates were transfected with 0.1 µg of plasmid DNA per well using LyoVec transfection reagent (InvivoGen), according to manufacturer instructions, incubated for 18 h at 37°C, fixed with 4% paraformaldehyde in PBS, and imaged using a Zeiss Axiovert 200 inverted microscope equipped with an HBO 100 mercury arc lamp or a Nikon STORM with A Hamamatsu ORCA Flash 4.0 CMOS monochrome camera and a Nikon DS-Ri2 color camera. For transfections with plasmids expressing untagged NSP1-1 proteins, F-actin was detected with rhodamine phalloidin, and nuclei were detected using 300 nM 4′,6-diamidino-2-phenylindole (DAPI, Invitrogen), with washes in PBS. For transfections with plasmids expressing tagged NSP1-1 proteins, cells were fixed with cold methanol at 18 h post transfection and blocked with PBS containing 1% FBS. Then, FLAG peptides were detected with mouse anti-FLAG diluted 1:500, and Alexa Fluor 546-conjugated anti-mouse IgG, diluted 1:1000, and nuclei were detected using 300 nM DAPI, with washes in PBS containing 0.5% Triton X-100.

Cos7 cells (∼6 × 10^4^ per well) in 24-well plates were transfected with 0.5 µg of plasmid DNA per well using LyoVec, according to manufacturer instructions, and incubated at 37°C. At 18 h post transfection, cells were fixed with 4% paraformaldehyde in PBS. Staining to detect F-actin and nuclei and imaging were conducted as described above.

BHK-T7 cells (∼6 × 10^4^ per well) in 24-well plates were transfected with 0.5 µg of plasmid DNA per well using TransIT-LT1 transfection reagent (Mirus Bio) in OptiMEM (Gibco), according to manufacturer instructions, and incubated at 37°C. At 18 h post transfection, cells were fixed with 4% paraformaldehyde in PBS. Staining to detect F-actin and nuclei and imaging were conducted as described above.

LLC-PK1 cells (∼1 × 10^5^ per well) in 24-well plates were transfected with 0.1 µg of plasmid DNA per well using LyoVec, according to manufacturer instructions, and incubated at 37°C. At 18 h post transfection, cells were fixed with 4% paraformaldehyde in PBS. Staining to detect F-actin and nuclei and imaging were conducted as described above.

DF-1 cells (∼1 × 10^5^ per well) in 24-well plates were transfected with 0.5 µg of plasmid DNA per well using LyoVec, according to manufacturer instructions, and incubated at 37°C. At 18 h post transfection, cells were fixed with 4% paraformaldehyde in PBS. Staining to detect F-actin and nuclei and imaging were conducted as described above.

MDCK.1 cells (∼6 × 10^4^ per well) in 24-well plates were transfected with 0.5 µg of plasmid DNA per well using LyoVec, according to manufacturer instructions, and incubated at 37°C. At 18 h post transfection, cells were fixed with 4% paraformaldehyde in PBS. Staining to detect F-actin and nuclei and imaging were conducted as described above.

### Quantitation of Syncytia Diameter

293T cells in 24-well plates were transfected with 0.1 µg per well of plasmids encoding untagged or tagged forms of RVB, RVG, or RVI NSP1 or control plasmids and stained to detect actin and nuclei (untagged) or FLAG and nuclei (tagged) as described for imaging studies. Syncytia were identified visually and imaged using a Zeiss Axiovert 200 inverted microscope. When possible, the person analyzing images was blinded to the identity of the samples. Diameters of at least 30 syncytia imaged from a minimum of four independently transfected wells of 293T cells per plasmid construct were quantified as the average of two diameter measurements per syncytium made using the measure function in Fiji (86). Statistical analyses of cluster diameters were conducted using Kruskal-Wallis tests with Dunn’s multiple comparisons in GraphPad Prism 9 (GraphPad).

## ACKNOWLEDGEMENTS

We thank the staff at the Vanderbilt Cell Imaging Shared Resource for assistance with image acquisition and analysis. We thank Dr. Takeshi Kobayashi for NBV MB p10 in pCAGGS and Dr. Marco Morelli for the pLIC6 plasmid. We are grateful to Alejandra Flores and Sydni Caet Smith for critical reading of the manuscript.

Research funding was provided by the National Institutes of Health (1R21AI153769). Cell Imaging Core Services performed through Vanderbilt University Medical Center’s Digestive Disease Research Center were supported by NIH grant P30DK058404. The contents of this publication are solely the responsibility of the authors and do not necessarily represent the views of the National Institutes of Health.

